# Fish Playpens – Method for raising individual juvenile zebrafish on a recirculating system for studies requiring repeated measures

**DOI:** 10.1101/2023.11.20.567840

**Authors:** Tabea O.C. Moll, Steven A. Farber

**Affiliations:** Johns Hopkins University, Baltimore, Maryland, United States of America

## Abstract

Even though many experimental approaches benefit from tracking individual larval animals, there is yet to be a commercial zebrafish rack system designed to accomplish this task. Thus, we invented playpens, an acrylic and screen container, to raise 12 individual zebrafish juveniles per standard 10 L tank on an existing recirculating fish system. During a week-long experiment, fish raised in playpens grow to the same size as conventionally raised juveniles.

---

Repeated assays of individual larvae are readily accomplished during the first 5 days of development when the larva relies entirely on the maternally deposited yolk as its food source^1^, in that they can be maintained in multi-well dishes. However, after 5 days post fertilization (dpf), there is no reliable way to raise individual juveniles on a recirculating zebrafish system. There are many protocols for raising zebrafish for research purposes, but they vary greatly, and none describe protocols to raise single juvenile animals. Original rearing guidelines described feeding larvae in beakers without water flow until 12 (day post fertilization) dpf before putting them on a recirculating or flow-through system^2,3^. However, this approach requires meticulous cleaning and care, is time-consuming, and can produce variable results. Because of these and other issues, many zebrafish facilities move larvae into tanks at 5 - 6 dpf with a gentle low water flow^4^. This way, 10 larvae per liter can be raised simultaneously, with considerably less feeding effort and better viability than most off-system methods^5^.

In this rearing method, time course studies require the use of a cross-sectional design and collection of different larvae at each time point. However, repeated measures of the same animal are often superior to a cross-sectional experimental design because they can better assay the impact of an intervention on one individual across time. In a repeated measures experimental design, each animal can serve as its own control, which results in the detection of smaller changes with higher statistical power^6^. For these reasons, a repeated measures experimental design is preferable to a cross-sectional approach, especially when investigating the effects of diet or drugs on digestive processes over time^7^. The Farber lab has developed various optical reporters to follow metabolic processes without euthanizing the animal^8-11^, a technology ideal for a repeated measures approach.

Thus, we developed a method to raise individual larvae in custom-designed “playpens” inserted into standard tanks (Fig. 1 A). This approach can be readily modified to be employed in any commercially available group housing tank. A playpen is constructed by taking a square acrylic tube (McMaster-Carr, 8516K41) and removing 3 of the 4 sides. The remaining side provides the needed structure to support the top and bottom 1.2 cm frame (Fig. 1 A). The open sides and bottom frame are closed off by a 400-micron screen (Dynamic Aqua Supply Ltd., NTX400-106), secured to the acrylic structure. Silicone (Loctite, 908570) is used to seal the screen and allowed to dry for 24 hours before use. This creates a square tube structure (12.8 cm high) with 3 screen sides and a screen bottom to contain a juvenile zebrafish while allowing for water and waste flow (Fig. 1 A, E). A standard 10 L tank was fitted with a false bottom (Home Depot, #1199233A) that supports the playpens at the correct height in the tank (4 cm high false bottom). Its grid structure with large holes allows debris and feces to fall from each playpen to the bottom of the tank and then be flushed from the tank. The upper square frame rises above the water level, allowing water circulation through the entire playpen (Fig. 1 B). Up to 12 playpens can be placed in a 10 L tank on the false bottom, each playpen housing one individual larva.

**Figure 1:**
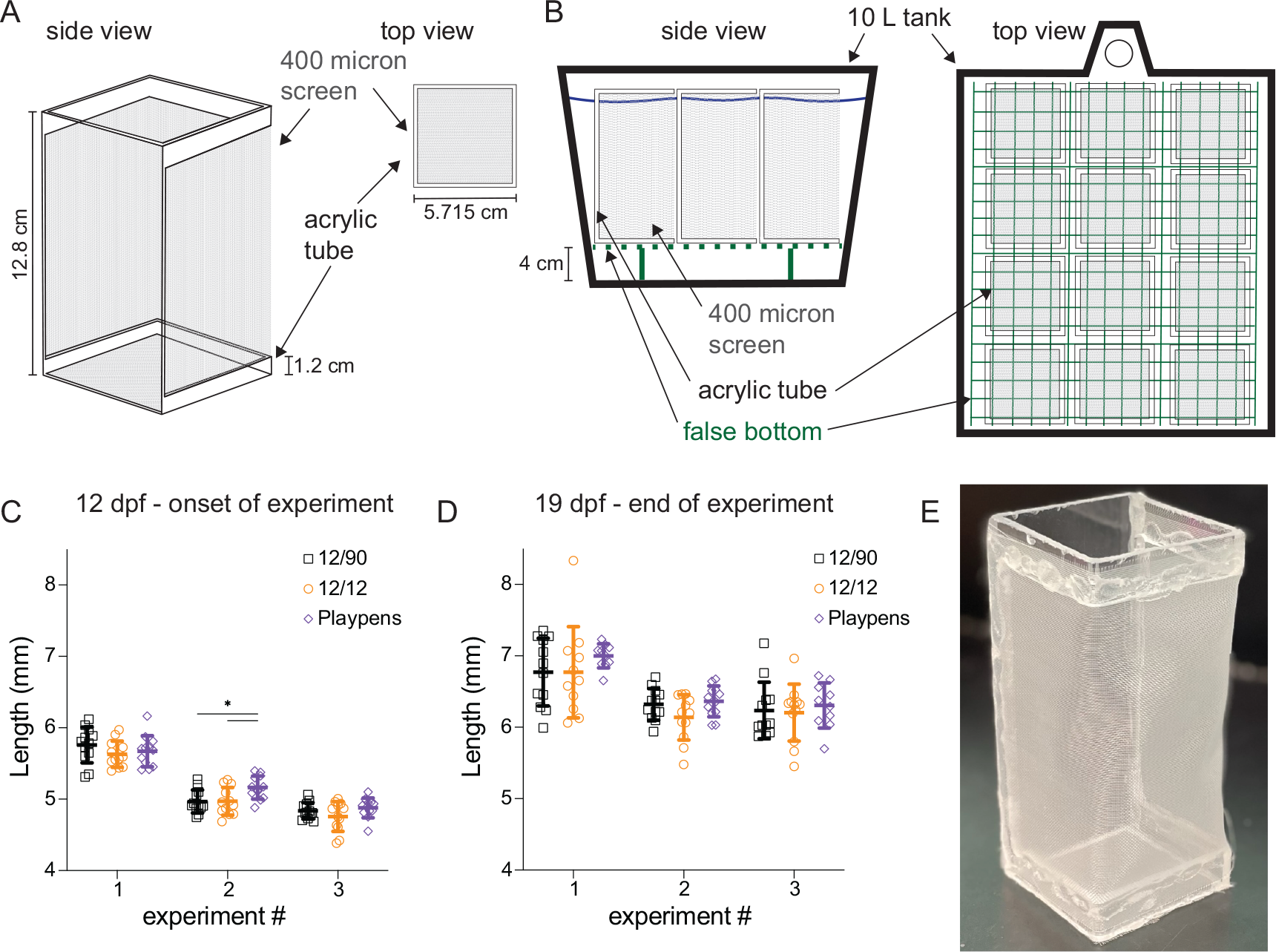
Playpens to raise individual zebrafish on fish nurseries lead to comparable growth of juveniles. Schematic of playpens to raise individual zebrafish in 10 L tanks on the nursery of a regular fish facility. (A) side and top view of an individual playpen. Three of the four sides of the acrylic square tubes are removed and replaced with 400-micron screen. An additional screen is placed on the bottom for the structure, preventing the fish from escaping through the bottom. The mesh is held in place using silicone. (B) 12 playpens can be placed on a false bottom, and the top square of the playpen is above the water line. This allows excess food and feces to fall through and be flushed out instead of trapped in the playpen. The top view shows the placement of 12 playpens on top of the false bottom in the 10 L tank. (C) At 12 dpf, 12 fish for each condition were measured. The fish of the playpens and the low-density housing (12/10L) were raised together and randomly chosen for further placement in the playpens or low-density setup. 12 fish from a normal 10L at 90/10L were measured and placed back into the tank. (D) After 1 week, the fish were measured again. No difference in size is detectable between the different housing conditions (2-way ANOVA followed by Tukey’s multiple comparisons test). (E) Photograph of one playpen.

We examined the growth of 12 larvae raised in three different conditions: in playpens; at low density by raising 12 larvae in a 10 L at the same overall density per liter as the playpens (we refer to as 12/10L); under standard conditions with 90 larvae in a 10 L tank (we refer to as 90/10L). This experiment was performed three independent times. Before the onset of each experiment, the juveniles for playpens and 12/10L were raised together in 10 L at the standard 90 larvae / 10 L density of our facility and then chosen randomly for the two groups. The length of larvae of each condition was measured at the beginning, at 12 dpf, and at the end of the housing experiment, at 19 dpf. For accurate size measurements, zebrafish were removed from the playpens using plastic pipettes (Amazon, B00G6T8OYS), mounted in 3 % methylcellulose (Sigma, M0387), and imaged with the Olympus Axiozoom V16 microscope equipped with a Zeiss Plan NeoFluar Z 1x/0.25 FWD 56 mm objective, AxioCam MRm camera, and Zen 2.5 software at 16 x magnification. The length of the fish was measured using FIJI and analyzed by GraphPad Prism (2-way ANOVA and Tukey’s multiple comparisons test).

The second time the experiment was performed (experiment 2), the randomly selected 12 dpf juveniles from the playpens were larger than the other two conditions (5.2 ± 1.6 mm in playpens compared to 5.0 ± 1.6 mm in 90/10L, p = 0.029 and 5.0 ± 1.9 mm in 12/10L, p = 0.033; Fig. 1 C). There was no significant difference in size at 19 dpf between any of the conditions at any experiment (Fig. 1 D), suggesting that raising juvenile zebrafish in playpens does not affect their growth rate. It is not surprising to see that overall fish growth varied between the three independent experiments since several external factors of a fish facility can lead to variations in juvenile growth rates that eventually stabilize in equal adult sizes.

In summary, this new method of raising individual juvenile zebrafish on the nursery system of a fish facility is of great interest when performing experiments that require uniform growth and/or the tracking of individual fish.

## Acknowledgments

We greatly thank Theodore J. Cooper for realizing our ideas of zebrafish playpens and his expertise in constructing them. We want to thank Mackenzie Klemek for helping with the assembly of the playpens and for feeding our zebrafish.

## Authorship Contribution

T.O.C.M.: conceptualization, investigation, methodology, writing – original draft. S.A.F.: Supervision, writing – reviewing and editing,

